# Temporal genomics help in deciphering neutral and adaptive patterns in the contemporary evolution of kelp populations

**DOI:** 10.1101/2023.05.22.541724

**Authors:** Lauric Reynes, Louise Fouqueau, Didier Aurelle, Stéphane Mauger, Christophe Destombe, Myriam Valero

## Abstract

The long-term persistence of species in the face of climate change can be evaluated by examining the interplay between selection and genetic drift in the contemporary evolution of populations. In this study, we focused on spatial and temporal genetic variation in four populations of the cold-water kelp *Laminaria digitata* using thousands of SNPs (ddRAD-seq). These populations were sampled from the center to the south margin in the North Atlantic at two different time points, spanning at least two generations. By conducting genome scans for local adaptation from a single time point, we successfully identified candidate loci that exhibited clinal variation, closely aligned with the latitudinal changes in temperature. This finding suggests that temperature may drive the adaptive response of kelp populations, although other factors, such as the species’ demographic history should be considered. Furthermore, we provided compelling evidence of positive selection through the examination of allele frequency changes over time, offering additional insights into the impact of genetic drift. Specifically, we detected candidate loci exhibiting temporal differentiation that surpassed the levels typically attributed to genetic drift at the south margin, confirmed through simulations. This finding was in sharp contrast with the lack of detection of outlier loci based on temporal differentiation in a population from the North Sea, exhibiting low levels of genetic diversity, that further decreased over time. These contrasting evolutionary scenarios among populations can be primarily attributed to the differential prevalence of selection relative to genetic drift. In conclusion, our study highlights the potential of temporal genomics to gain deeper insights into the contemporary evolution of marine foundation species in response to rapid environmental changes.

## 1. INTRODUCTION

The different selective pressures that arise from varying environments, both spatially and temporally, can lead to distinct trait values through local adaptation, phenotypic plasticity, or a combination of both (de Villemereuil et al., 2018; Reger et al., 2018). When population divergence is driven by adaptation, gene flow between populations can serve as an important source of genetic variation for traits under selection. The rate at which populations respond to environmental change is directly proportional to the genetic variation present in fitness-related traits, particularly when adaptation arises from existing genetic variation (Barrett & Schluter, 2008). Local adaptation is thus considered a promising phenomenon that can facilitate species’adaptation to climate change (Aitken et al., 2008; Savolainen et al., 2013). However, genetic differentiation also arises from stochastic processes, such as genetic drift, especially in small and isolated populations. These populations, with limited gene flow, may experience reduced genetic diversity, which can impede or slow down the process of local adaptation (Lande, 1988; Willi et al., 2006). Therefore, it is important to disentangle the effects of natural selection from neutral processes for assessing the adaptive potential of natural populations in the face of rapid environmental change. This knowledge holds value in the development of genetically informed ecological niche models (Ikeda et al., 2017; DeMarche et al., 2019) and the implementation of effective conservation strategies that integrate adaptive patterns of genetic variation and assess their potential mismatch with future climate change (e.g., Sgrò et al., 2011; Von der Heyden, 2017; Xuereb et al., 2020).

Genome-wide screening of genetic variation has provided evidence supporting the potential for populations to undergo local adaptation to specific environmental conditions, despite experiencing substantial rates of genetic drift. Indeed, several studies have identified candidate loci for positive selection in relatively small, structured populations (e.g., Funk et al., 2016; Perrier et al., 2017; Leal et al., 2021; Pratlong et al., 2021), raising questions regarding the relative contribution of selection and genetic drift. However, it is crucial to acknowledge the potential impact of factors, such as high variance in F_ST_ among loci and other sources of spatial variation in allele frequencies on the sensitivity of genome scans, particularly those utilizing F_ST_-based approaches (Storz, 2005; Bierne et al., 2011; Lotterhos & Whitlock, 2015; Hoban et al., 2016). Furthermore, the effectiveness of these approaches can be limited when detecting weak or recent events of positive selection or when selection acts upon numerous loci with small effect sizes (de Villemereuil et al., 2014).

One potential improvement to enhance genome scans for local adaptation is the inclusion of samples collected at multiple time points (see for review Clark et al., 2023). This approach offers valuable insights into both neutral and adaptive patterns of genomic variation, as demonstrated by studies, such as Therkildsen et al. (2013a, 2013b) and Frachon et al. (2017). The underlying premise of temporal genomics is that loci under positive selection exhibit a consistent directional trend in allele frequency, which allows them to be distinguished from random changes due to genetic drift. Hence, comparing temporal genetic differentiation expected by genetic drift using forward genetic simulation (Goldringer & Bataillon, 2004), as well as, assessing, covariance in allele frequencies over time (Buffalo & Coop, 2019, 2020), can be informative for detecting loci contributing to temporal genetic changes underlying rapid adaptation to novel selective pressures. Temporal genomics also provides an opportunity to measure temporal variance in allele frequencies, which is expected to provide insight into contemporary estimates of effective size (Ne) compared to methods using linkage disequilibrium (Waples, 1989, 2005). This has significantly enhanced the power to detect declines and more accurate inferences of Ne, particularly when historical population sizes were large (Nadachowska-Brzyska et al., 2021; Reid & Pinsky, 2022).

The Laminariales, commonly known as kelps, present an interesting model for investigating the interplay between selection and genetic drift in the contemporary evolution of populations to rapid climate warming. Understanding the ability of such cold-temperate water species to adapt to warmer conditions is crucial in assessing their vulnerability to local extinction (Araújo et al., 2016; Arafeh-Dalmau et al., 2019; Filbee-Dexter et al., 2020). Recent studies on *Laminaria digitata* in the Northern Hemisphere have suggested that populations at the south margin may exhibit lower sensitivity to heat stress compared to those in the North (Liesner et al., 2020; Schimpf et al., 2022). Although significant progress has been made in understanding the genetic basis of thermal preference and heat tolerance in kelp species (Mao et al., 2020; Guzinski et al., 2020; Vranken et al., 2021), identifying candidate loci by genome scans for local adaptation at a single time point remains challenging due to pronounced levels of neutral differentiation. Kelps typically display considerable genetic structuring attributed to limited dispersal capacities and seascape features, leading to relatively low to moderate effective population sizes at the trailing edge (Coleman et al., 2011; Brennan et al., 2014; Assis et al., 2022; Fouqueau et al., 2023). Based on the current state of knowledge, the question of whether populations at the southern margin are locally adapted to warmer temperatures or if the previously observed patterns of thermal adaptation arose from phenotypic plasticity remains unresolved.

The main objective of our study was to investigate the impact of local selection and genetic drift on the contemporary evolution of the kelp *Laminaria digitata* across both spatial and temporal dimensions. To achieve this, we employed a reduced representation sequencing technique (ddRAD-seq, Peterson et al., 2012) on four populations, with three of these populations sampled at two different time points. Our specific aims were as follows: i) estimating contemporary effective population size (Ne) based on temporal genetic changes to assess whether limited genetic variation at the southernmost margin may hinder the population’s adaptive response, ii) conducting genome scans across space to identify potential signatures of selection in response to thermal conditions, iii) differentiating the effects of genetic drift from selection by analyzing changes in allele frequencies over time, allowing us to detect potential signatures of ongoing selection or genetic erosion that may occur within a few generations. Overall, our results point out the promise of temporal genomics to gain deeper insights into the contemporary response of this fundamental marine species to rapid environmental changes.

## 2. MATERIAL AND METHODS

### 2.1 Sampling

Populations of *Laminaria digitata* were sampled from four locations spanning from the species’ center to its southern margin. The sampling sites included Clachan, Scotland (CLA); Helgoland, Germany (HLG); Roscoff, France (ROS) and Quiberon, France (QUI) (Table 1, Figure 1). Tissue samples of approximately 3cm were collected from 30-40 sporophytes in the low intertidal zone at each site. The samples were promptly stored in silica gel to preserve their genetic material. For three out of the four populations (HLG, ROS, QUI), samples were collected at two different time points to capture changes in allele frequencies over a short period and gain insights into the relative influence of directional selection and genetic drift. The time interval between the two sampling periods ensured that at least two generations of sporophytes were included in the study (Bartsch et al., 2008). The first set of samples was collected between 2005 and 2011, while the second set was collected in 2018. Regarding the CLA population, samples were collected in 2008 and 2018 from two separate sites, CLA_1 and CLA_2, which were approximately 20 km apart. Given the frequent observation of genetic structure within the species at scales of 1-10 km (e.g., Billot et al., 2003, Robuchon et al., 2014), the samples from CLA were treated as separate populations rather than multiple time points of the same population.

**Table 1.** Geographical coordinates and genetic diversity of populations at two different time points.

| Population | Year | Lat. | Lon. | n | P (± SE) | He (± SE) |
| --- | --- | --- | --- | --- | --- | --- |
| Helgoland (HLG) | 2005 | 54.178 N | 7.893 E | 18 | 0.11 (0.001) | 0.015 (0.001) |
|  | 2018 |  |  | 22 | 0.08 (0.001) | 0.013 (0.001) |
| Clachan (CLA_1) | 2008 | 56.454 N | 5.444 W | 25 | 0.42 (0.001) | 0.103 (0.003) |
| Clachan (CLA_2) | 2018 | 56.318 N | 5.592 W | 25 | 0.42 (0.001) | 0.109 (0.003) |
| Perharidy (ROS) | 2011 | 48.727 N | 4.005 W | 27 | 0.36 (0.001) | 0.065 (0.002) |
|  | 2018 |  |  | 25 | 0.36 (0.001) | 0.069 (0.002) |
| Quiberon (QUI) | 2009 | 47.470 N | 3.091 W | 24 | 0.21 (0.001) | 0.038 (0.001) |
|  | 2018 |  |  | 24 | 0.20 (0.001) | 0.040 (0.002) |

Year: the specific year when the samples were collected; Lat: latitude coordinate of the sampling location; Lon: longitude coordinate of the sampling location; n: the number of individuals that were genotyped; P ( ± SE): the proportion of polymorphic loci using a minor allele frequency threshold of 0.01 (mean and ± SE); He ( ± SE): expected heterozygosity, represented as the mean value with standard error (SE)

**Figure 1.**
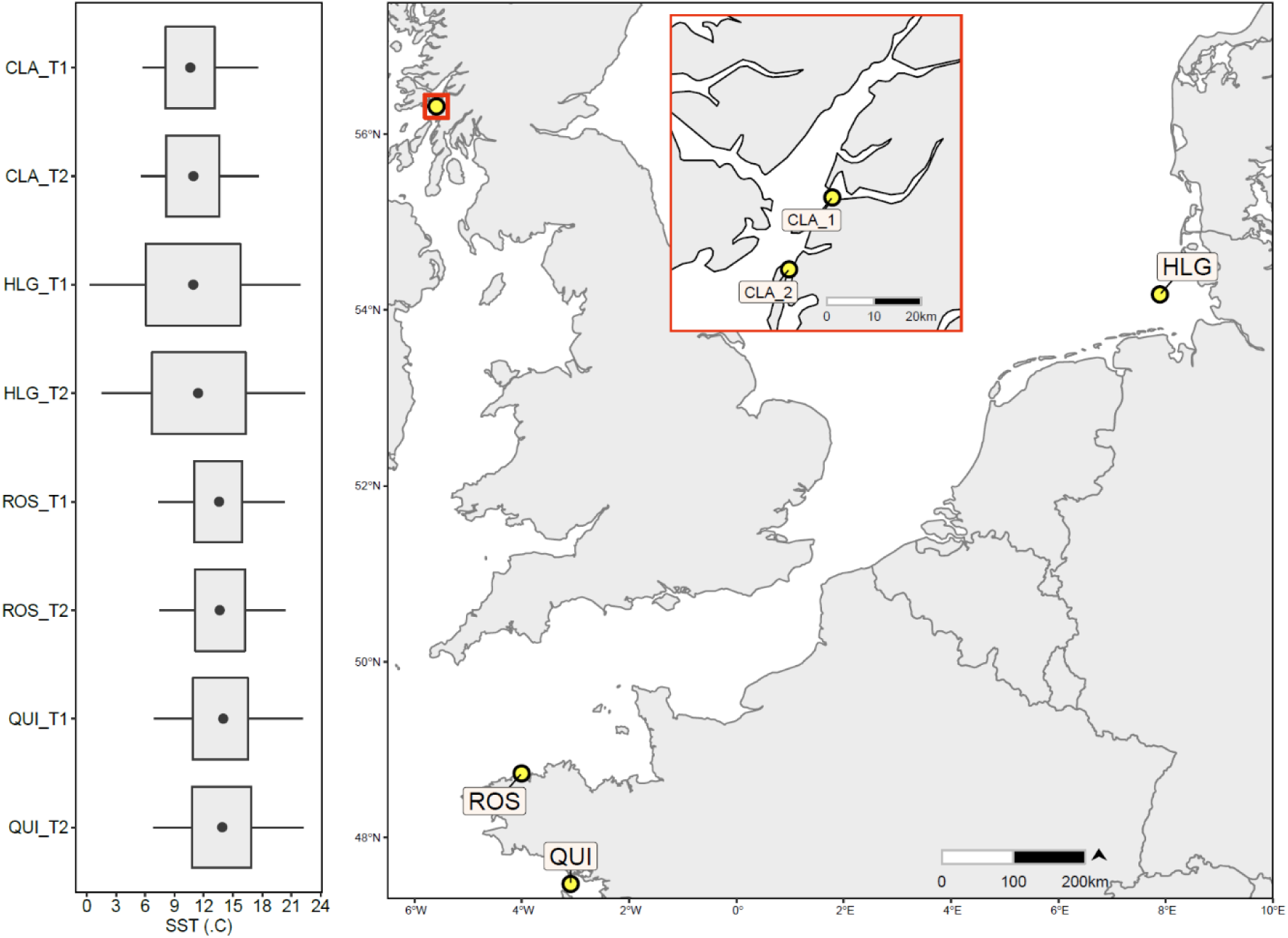
Sampling sites of the *Laminaria digitata* populations used in this study. Sea Surface Temperature (SST) was represented as a boxplot depicting variation at a focal site during both time points (T1 and T2).

### 2.2 Temperature data acquisition

Given the increasing body of evidence supporting the role of temperature in driving adaptation within the species (King et al., 2019; Liesner et al., 2020; Schimpf et al., 2022) sea surface temperature (SST) was considered the most relevant factor for investigating adaptive differentiation among populations. To characterize the SST ranges experienced by the populations, we utilized daily mean SST data obtained from satellite observations with a resolution of 0.05° × 0.05°, sourced from the E.U. Copernicus Marine Service Information (E.U. Copernicus Marine Service, 2022). In order to account for the influence of ongoing climate warming on temporal genetic changes, we differentiated SST predictors between two distinct periods: 1993-2000 and 2000-2018. This approach allowed us to consider the slight increase in SST experienced by the respective generations during the first and second time points.

### 2.3 Library preparation, SNP and genotype calling

DNA extraction was performed on dried tissue that has been preserved in silica gel for several years using the Nucleospin R 96 plant kit II (Macherey-Nagel, Düren, Germany), following the manufacturer’s protocol. Two double-digest RAD-sequencing libraries (ddRAD-seq, Peterson et al., 2012) were prepared, with 117 individuals for the first time point and 104 individuals for the second time point, including eight replicates (i.e. eight samples with the same DNA extract but independent library preparation, sequencing, reads mapping, and SNP calling) in each population and time point. Individuals were randomly distributed across the libraries, along with 355 samples of distinct projects to prevent batch effects and ensure library diversity. Library preparation was conducted according to Reynes et al., (2021), using 100 ng of DNA and the *PstI* and *HhaI* enzymes (NEB). Paired-end reads with 150 cycles were obtained by sequencing the libraries on an Illumina Hiseq 4000 platform (Génome Québec Innovation Centre, McGill Univ., Montreal, Canada). Quality control of the sequences was performed using FASTQC v.0.11.7 (Andrews, 2010). Adaptors were removed, and the reads were trimmed to 137 bp using Trimmomatic (Bolger et al., 2014). The paired-end reads were then mapped to the draft reference genome of *L. digitata* using BWA-mem with default parameters in BWA v.0.7.17 (Li & Durbin, 2009). The N50 of the draft genome is ∼9.9 Mb with a genome assembly size of ∼470 Mb among 127 552 scaffolds. Uniquely mapped reads were retained with SAMtools v.1.13 (Danecek et al., 2021). SNP calling was performed with the reference mode of the Stacks v.2.52 pipeline (Catchen et al., 2011). Genotyping and SNP calling were carried out separately for individuals at each time point, and the shared SNPs between the datasets were selected before merging individuals across these SNP positions. The *bcftools isec* and *bcftools merge* functions of BCFtools v.1.9 (Danecek et al., 2021) were used for these steps. Post-call filtering of SNPs was performed on the merged dataset by keeping SNPs with a call rate >90% per population in one or more populations using *pop_missing_filter.sh* of the dDocent pipeline (Puritz et al., 2014). Problematic individuals (n = 23) with a call rate below 80% were discarded. Filtering based on mean read depth (DP) and minor allele frequency was implemented using vcftools v.0.1.16 (Danecek et al., 2011), with specific parameters outlined in Table 2. To address the effects of excessive linkage disequilibrium (LD) between loci, SNPs with a square correlation (r^2^) >0.2 were removed using PLINK v.1.9 (Chang et al., 2015). The calculation of r^2^ values was performed separately for each population to separate the effects of physical proximity among SNPs from the effects of genetic structure in LD patterns. SNPs with r^2^ values exceeding the threshold in all populations were discarded (Table 2). After all post-filtering steps, a total of 2 854 SNPs remained among 190 individuals (excluding the eight replicates). Genotyping concordance was assessed in this dataset by calculating the SNP error rate between replicate pairs following Mastretta-Yanes et al., (2015).

**Table 2.** The filtering steps applied to the raw sets of 77 193 SNPs and 644 794 SNPs for the first and second time points, respectively.

| Filtering steps | SNP | Individual |
| --- | --- | --- |
| SNPs present in both the first and second time points | 36 293 | 213 |
| Shared SNPs among populations with less than 10% of missingness per SNP within a population | 4 151 | - |
| Individual missingness rate $\leq 0.2$ | 4 151 | 190 |
| Mean read depth (DP); SNPs between 10x and twice the mean. For the first time point, the mean read depth was 34x, and for the second time point, it was 45x | 3 995 | - |
| SNPs deviating from HWE ($P < 0.001$) in at least 25% of the populations | 3 995 | - |
| Minimum allele count of three | 3 609 | - |
| SNPs pruned at an LD threshold of $r^2 = 0.20$ within and among populations | 2 854 | - |

### 2.4 Genetic differentiation

The genetic differentiation among populations and over time was assessed using an analysis of molecular variance (AMOVA) implemented in Arlequin v.3.11 (Excoffier et al., 2005), with 10 000 permutations. To account for temporal changes, the AMOVA considered genetic variation between two time points within each of the three populations that were resampled over time. Spatial and temporal genetic differentiation were quantified by calculating F_ST_ (Weir and Cockerham, 1984) using the R package diveRsity v1.9.90. The significance of F_ST_ values was evaluated using an exact test of genic differentiation (Raymond and Rousset, 1995) with the Markov chain method and default parameters. A combination of all tests across loci was performed using Fisher’s method. Genetic structure was investigated using the sNMF algorithm implemented in the R package LEA v.2.8 (Frichot et al.,2014; Frichot and François, 2015). The analyses involved 10 000 iterations and 20 repetitions, with K ranging from 1 to 16. The best K value was determined based on the cross-entropy criterion. A complementary analysis of genetic structure was performed using a principal component analysis (PCA) on SNP variation, utilizing the R packages Bigsnpr v.1.3.0 and Bigstatsr v.1.2.3 (Privé et al., 2018). Missing genotypes were imputed by replacing them with the average allele frequencies before performing the PCA.

### 2.5 Genetic diversity and effective population size

Genetic diversity within populations was assessed using the complete set of 2 854 SNPs. Expected heterozygosity (He) was estimated using the R package diveRsity v.1.9.90. Differences in He between populations and time points were evaluated using pairwise Wilcoxon tests, with adjustment for multiple comparisons using the Bonferroni method. To estimate the proportion of polymorphic loci (P%) within each population and time point, a custom bash script (available upon request) was used. The P% was calculated by randomly sampling 20 individuals (with replacement) and iterating the procedure 100 times. A minor allele frequency threshold of 0.01 was applied, and averages and standard errors of P% were computed. The contemporary effective population size (Ne) was estimated based on the temporal variance in allele frequencies using three methods: two methods based on F-statistics Fc (Nei & Tajima, 1981) and Fs (Jorde & Ryman, 2007) and a likelihood estimator (Wang & Whitlock 2003), known for its improved precision and accuracy in the presence of rare alleles (Wang, 2001). For the F-statistics methods, the software Neestimator v.2.1 (Do et al., 2014) was employed, while the likelihood approach was performed using MLNe v.2.0 (Wang & Whitlock, 2003). The 95% confidence intervals of the likelihood estimate and moment F statistics were also calculated. The Ne estimation considered the plan II sampling procedure, assuming individuals are sampled before reproducing and are not returned to the population (Waples, 1989). Additionally, it was assumed that two generations had passed between the time points.

### 2.6 Outlier detection across space

Outlier tests based on spatial differentiation were conducted to identify loci that exhibited patterns of differentiation among populations deviating from neutral expectations, potentially indicating the influence of directional selection. These outlier tests were performed separately for each time point. Individuals from CLA_1 (2008) and CLA_2 (2018) were included in these tests, despite their exclusion from analyses based on temporal genetic changes. Three different methods were employed for outlier detection across space. Firstly, the Bayesian approach of Beaumont & Balding (2004), implemented in BayeScan 2.1 (Foll & Gaggiotti, 2008) was used. The program was run with different prior odds (3, 5, 10 and 100) with 20 pilot runs of 5 000 iterations each, followed by a burn-in of 50 000 iterations and 5 000 samplings. The second method utilized a principal component analysis implemented in the R package pcadapt (Luu et al., 2017). Pcadapt assessed the contribution of individual SNPs to K principal components, considering their relationship to population structure. Lastly, OutFLANK v.0.2 software (Whitlock & Lotterhos, 2015) was employed, which identifies outliers by comparing differentiation at each SNP against a trimmed null distribution of F_ST_ values. OutFLANK was run with LeftTrimFraction = 0.55 and RightTrimFraction = 0.10. To account for multiple testing, the p-values obtained from each method were corrected using the R package qvalue v.2.18, with a threshold set at 0.10. Overlapping outlier SNPs identified by multiple methods and datasets were analyzed using jvenn (Bardou et al., 2014) to assess the common outliers across the different approaches.

### 2.7 Outlier detection across time

Outlier tests were performed between time points to differentiate the effects of selection from genetic drift in allele frequency changes. Initially, using SLiM 3 (Haller & Messer, 2019), we simulated SNP frequencies 5000 times to evaluate the expected level of genetic differentiation after two generations of genetic drift alone. The initial SNP frequencies for each simulation matched the frequencies observed in our samples at the first time point. We assumed panmictic reproduction while neglecting mutation and migration. N individuals were sampled, considering the previously estimated Ne (see Table 3), to establish the next generation. Genetic differentiation under genetic drift was then evaluated after two generations by sampling n individuals of the same size as the second time point’s sample. This framework allowed for the detection of candidate loci undergoing selection, which exhibited higher temporal differentiation than expected from neutral simulations. For each of the three populations considered (HLG, ROS, QUI), simulations were conducted using the lowest and highest estimates of Ne based on temporal methods. Subsequently, we assessed the effect of Ne on the detection of outlier SNPs (see Supplementary S1). The vcf output files, which included empirical and simulated SNPs (5000 files for each population and Ne scenario) were processed using vcftools v.0.1.16 to calculate pairwise F_ST_. Finally, the p-value of the outlier test, which corresponded to the proportion of simulated F_ST_ equal and larger than the observed F_ST_ was calculated at a focal SNP using a custom R script (available upon request). We compared the outlier test based on SLiM to the method implemented in TempoDiff, which aimed to distinguish neutral from selected polymorphisms, using temporal differentiation (Frachon et al., 2017). Additionally, we ran BayeScan, OutFLANK and pcadapt using the same parameters as reported in the “Outlier detection across space” of the manuscript. The p-values of the tests were corrected for multiple testing before conducting an overlap analysis among methods, as described in the above section.

**Table 3.** Contemporary effective population size (Ne) based on temporal genetic changes using two F-statistics: Fc (Nei & Tajima, 1981), Fs (Jorde & Ryman, 2007), and the likelihood estimator implemented in MLNe (Wang & Whitlock, 2003).

|  | HLG | ROS | QUI |
| --- | --- | --- | --- |
|  | Ne [IC] | Ne [IC] | Ne [IC] |
| $F_c$ | 59 [26 - 469] | 451 [157 - Inf] | 165 [77 - 3412] |
| $F_s$ | 45 [23 - 1403] | 247 [111 - Inf] | 104 [56 - 738] |
| likelihood | 104 [70 - 195] | 43 063 [935- Inf] | 227 [134 - 667] |

### 2.8 SNPs-temperature associations

To investigate SNPs potentially under selection, we examined whether their allele frequencies were correlated with Sea Surface Temperature (SST). We specifically focused on SNPs identified as outliers by at least two outlier tests across space. For these loci, we tested the association between SNPs-temperature following the logistic regression framework described by Bataillon et al. (2022). The primary objective was to identify SNPs whose frequencies were correlated with temperature while controlling for neutral patterns of genetic structure. To achieve this, we employed a binomial Generalized Linear Model (GLM) with log link (‘glm’ function in R). The response variable was the count of the reference allele at a focal SNP, coded as 0, 1, or 2. The model accounted for the effect of genetic structure by including the first five PCs that explained 68.1% of the variation of the PCA of SNP variation based on all SNPs. The assess the effect of temperature, we considered the first two PCs derived from the PCA conducted on daily mean SST statistics, including the average, minimum, maximum, and amplitude coefficient values (see Temperature data acquisition section). Only the first two PCs were retained as they accounted for 74.5% and 25.4% of the variation respectively, and effectively distinguished populations, likely reflecting SST differences between regions. Before performing the PCA, all SST predictors were centred and normalized. Three distinct GLMs were fitted: A background model that included PC1 to PC5 of SNP variation, and two additional models (ENV1 model and ENV2 model) that incorporated PC1 or PC2 of daily mean SST parameters as supplementary predictors of the five SNP variation predictors. Non-significant variables were initially removed using the ‘dropterm’ function from the R package MASS v.7.3. The model fit was evaluated using McFadden’s pseudo R-squared and compared to each other using the likelihood ratio test, implemented in the R package lmtest v.0.9. The p-values from the likelihood tests were adjusted using Bonferroni correction to account for multiple testing, as the association between SST and allele count was tested across multiple SNPs.

### 2.9 Gene ontology analysis

Candidate loci identified through outlier tests across space and time by at least two methods were subjected to functional annotation using ‘Omics Box’ v.1.3.11 (Götz et al., 2008). Initially, candidate loci were annotated using the NCBI Basic Local Alignment Search Tool (BLAST, Johnson et al., 2008) with the non-redundant protein sequences database. The BLASTx approach was employed, with a specific focus on Phaeophyceae sequences. The resulting BLAST hits were further mapped to the Gene using InterProScan (Zdobnov & Apweiler, 2001) and Gene Ontology (Ashburner et al., 2000) analysis was conducted. Finally, the results from both analyses were merged to obtain comprehensive functional annotation for the candidate loci.

## 3. RESULTS

### 3.1 Sequencing, SNP filtering and data quality

A total of 98.7 million reads were obtained for the first time point, while 327 million reads were obtained for the second time point. The individual proportion of mapped reads was on average 86.6% for the first time point and 95.5% for the second time point. Both the count of high-quality reads and the mapping rate were significantly lower in the first time point compared to the second time-point (Wilcoxon test, p-value < 0.001). As a result, the SNP discovery was approximately 8.3 times higher for the second time point (644 794 SNPs) compared to the first time point (77 193 SNPs). After filtering for shared loci between time points, with a call rate >90% per population in one or more populations, a total of 4 151 SNPs were retained (Table 2). Additional quality filtering, including individual missingness, read depth, minor allele frequency (MAF), and linkage disequilibrium further refined the dataset to a final set of 2 854 SNPs across 190 individuals, excluding eight technical replicates (Table 2). The analysis of eight technical indicated a high level of genotype concordance for the set of 2 854 SNPs, which was consistent across both time points. Specifically, the SNP error rate ranged from 0.007 to 0.036 for the first time point and from zero to 0.02 for the second time point (Table S1). The mean read depth was 16.73 X ± 17.93 SE for the first time point and 22.52 X ± 14.15 SE for the second time point. Despite the difference in sequencing depth between the two-time points, there was no significant difference in heterozygosity per individual (Wilcoxon test, p-value = 0.23).

### 3.2 Genome-wide genetic diversity

Overall, expected heterozygosity (He) showed significant differences between sites (Wilcoxon test, p-value < 0.001). the northern population (CLA_1 and CLA_2) exhibited the highest value of He, which was seven times higher than the population experiencing the most challenging summer temperatures for reproduction (HLG, Bartsch et al., 2013). Furthermore, the He values in CLA_1 and CLA_2 were 1.5 to 2.7 times higher compared to those observed in Brittany (ROS and QUI, respectively, Table 1). In the temporal comparison, it was found that He slightly decreased (∼2%) in HLG (Wilcoxon test over SNPs, p-value < 0.001), while its slight increase in Brittany was not statistically significant (Wilcoxon test over SNPs p-value = 0.13 for ROS and p-value = 0.047 for QUI). Moreover, a reduction (∼3%) in the proportion of polymorphic loci over time further supported the decrease in genetic diversity in HLG (Table 1). The contemporary estimation of Ne based on allele frequency variation over time for HLG, ROS and QUI aligned with the observed patterns in He and P%. Specifically, significant differences in Ne were reported between HLG (59-104) and QUI (104-227), in contrast to ROS (247-43 063). The different values for each population represented the range of Ne estimation obtained from F-statistics estimators and the likelihood-based estimator (Table 3).

### 3.3 Genetic structure

The AMOVA analyses conducted on the complete SNPs dataset revealed that variation was primarily attributable to variance within individuals (81%, p-value < 0.001) and between populations (21%, p-value=0.002). However, the effect of time was not significant (−0.61%, p-value = 0.708, Table 4). The average F_ST_ values among populations ranged from 0.196 to 0.221 for the first and second time point respectively. When excluding CLA_1 and CLA_2, the average F_ST_ values were 0.100 and 0.129 for the first and second time point respectively. All pairwise F_ST_ values in space were greater than or equal to 0.078 with a maximum value of 0.297 observed between CLA_2 and the second time point of HLG (Table S2). In space, all pairwise tests of genic differentiation showed significance (p-values < 0.001), while in time, they did not exhibit significance (p-values = 1), which was supported by temporal pairwise F_ST_ values close to zero (Table S2). The lack of significant differentiation over time using the complete SNPs dataset was further confirmed by the Sparse Non-negative Matrix Factorization (sNMF), as there was no significant change in genetic structure between time points (Figure S1). In contrast, there was a substantial genetic structure in space, with the northern populations (CLA_1 and CLA_2) showing differentiation from HLG ROS and QUI when considering K = 2. Results obtained at K = 3 differentiated ROS from QUI and HLG, and this was supported by the individual coordinates of these populations on PC1 and PC2 of the PCA of SNP variation (Figure S2). Finally, when considering the most informative number of clusters according to the minimum cross-entropy criterion (K = 5), individuals were grouped by sites, with an additional distinction between CLA_1 and CLA_2.

**Table 4.** Analyses of molecular variance (AMOVA) using the complete SNPs dataset

| Source of variation | D. | Var. components | Variation (%) | P-value |
| --- | --- | --- | --- | --- |
| Among populations | 4 | 16.07 | 20.71 | 0.00188 |
| Between time points | 3 | -0.47 | -0.61 | 0.70832 |
| Within populations | 182 | -0.88 | -1.14 | 0.91881 |
| Within individuals | 190 | 62.93 | 81.04 | < 0.00001 |

### 3.4 Outlier SNPs based on spatial differentiation

Among the 261 outliers (9.1 %) identified by at least one method, 97 (3.4%) were detected at both time points, indicating that genetic differentiation at these focal SNPs was conserved over time. On the other hand, 47 SNPs were exclusively detected in the first time point, and 88 SNPs were exclusively detected in the second time point. Interestingly, 36 SNPs were also identified as outliers by the temporal data-based tests. To reduce the number of false positives, we further considered only those SNPs that were detected by at least two out of the three detection methods. Consequently, no outlier SNPs remained for the first time point, while 16 SNPs remained for the second time point. Among these 16 SNPs, 15 were detected by both OutFlank and pcadapt, and only one SNP was detected by all three methods (OutFlank, pcadapt and BayeScan).

### 3.5 Outlier SNPs based on temporal differentiation

A total of 149 SNPs (5,2%) were identified by at least one method among the simulation framework described in the Methods section, TempoDiff and OutFLANK. BayeScan and pcadapt did not detect any outliers. The outlier tests based on our simulation framework reported the occurrence of 127 and 131 SNPs with higher F_ST_ than the neutral expectations when using, respectively, the lowest and highest Ne estimates. Due to the high correlation (r = 0.98) between p-values of the outlier tests conducted for the low and high Ne scenarios (Supplementary S1), we focused on the latter scenario, which utilized the Ne value estimated by the likelihood method (see Table 3). In this configuration, out of the 53 SNPs detected by TempoDiff, 35 (66%) were also identified by the simulation framework, including 13 SNPs that were detected by all three methods (simulations, TempoDiff and OutFLANK). When considering only the SNPs detected by at least two methods, the number of outlier SNPs decreased to 38, which were not detected by genome scans across space. Among these 38 SNPs, 23 were detected in ROS and 15 in QUI, while none were present in HLG (Figure 2A). Furthermore, we observed a substantial increase in observed heterozygosity (Ho) over time for these SNPs compared to putatively neutral ones. Specifically, Ho increased over time from 0.125 to 0.192 for ROS and from 0.087 to 0.184 for QUI. This pattern starkly contrasted with the low Ho values observed at HLG, which decreased from 0.033 to 0.021 over time. The increase in heterozygosity at QUI and ROS for these focal SNPs was strongly supported by high temporal F_ST_ values that exceeded 2.4 to 3 times the threshold of the 99% upper limit of neutral expectation, as determined by simulations (Figure 2BC, Figure 3). Conversely, only a very small number of SNPs exclusively detected by this method reached the 99% upper limit of neutral expectation at HLG.

**Figure 2.**
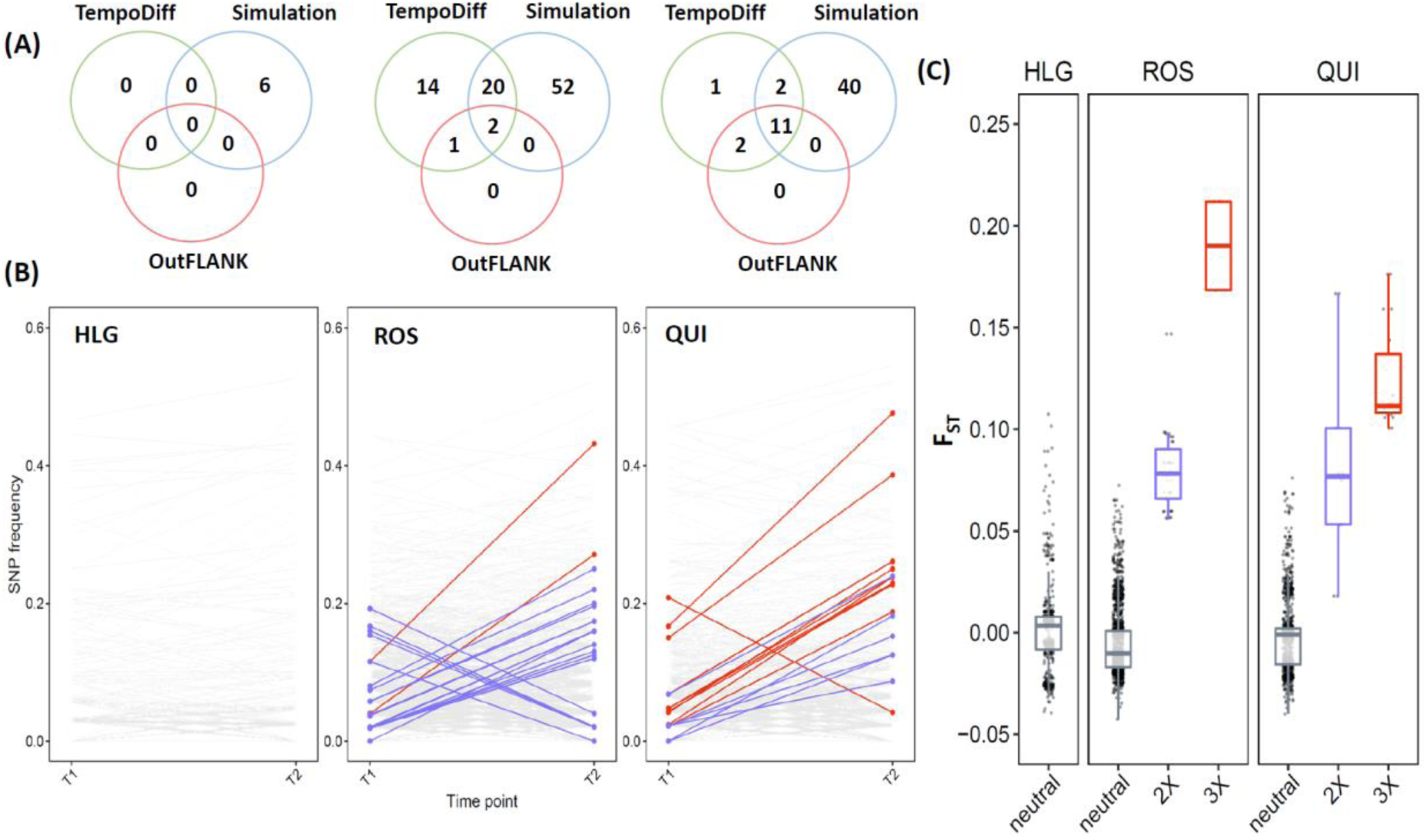
Outlier SNPs based on temporal differentiation are shown, depicting (A) the unique or shared outlier SNPs among methods for HLG, ROS, and QUI. The (B) allele frequency between time points and (C) F_ST_ values are provided depending on whether the SNPs were detected by none or only one method (grey), two methods (2X; purple) and three methods (3X; red). It is worth that the total number of SNPs in the Venn diagrams (A) is not equal to 149 SNPs, as two outliers at ROS were also detected at QUI by the simulation framework.

**Figure 3.**
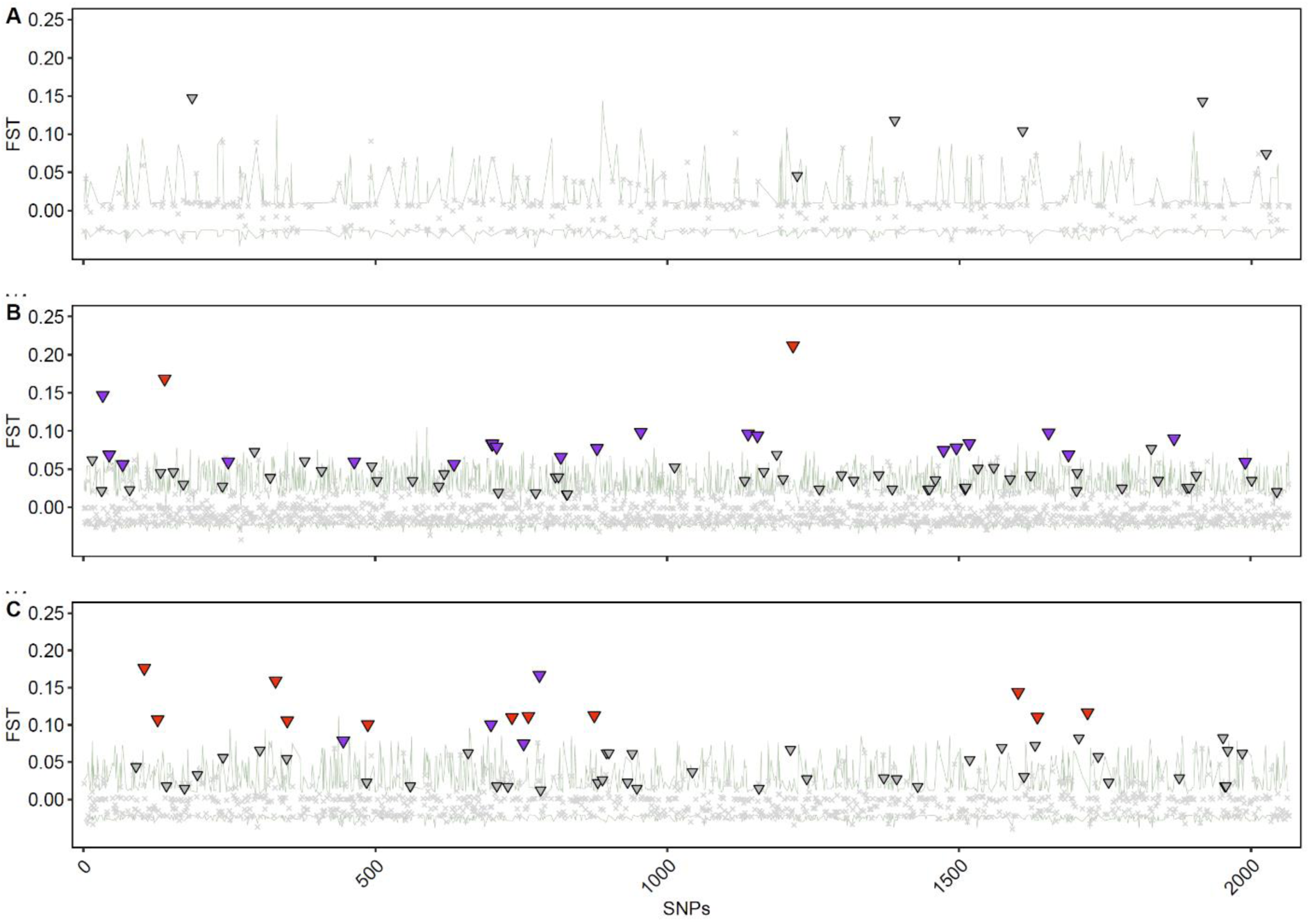
Temporal F_ST_ values of focal SNPs are compared to the upper limit of the neutral expectation based on the high Ne scenario. The green line represents the 99% quantile of F_ST_ values expected under genetic drift alone, determined from 5000 simulations for (A) HLG, (B) ROS, and (C) QUI. Outlier SNPs that are significant after correction for multiple comparisons and detected only by the simulation framework are represented by grey triangles. Those detected by two or three methods are shown in purple and red triangles, respectively. Grey crosses indicate putatively neutral SNPs that were not detected by outlier tests.

### 3.6 SNPs-temperature associations

We conducted association tests between the 16 outliers detected across space by at least two methods and SST using GLMs to account for the confounding effect of genetic drift in genetic structure. The results showed that incorporating PC1 or PC2 of the PCA performed on SST predictors, in addition to the five PCs of SNPs variation, resulted in a better fit for genetic differentiation (pseudo R^2^ = 0.50-0.84) for eight out of the 16 SNPs compared to the background model (pseudo R^2^ = 0.42-0.63; Table 5), which only included neutral predictors. However, the two SST-based PCs did not contribute equally to the patterns of genetic differentiation, as PC1 was never selected after performing one-term deletions (Table 5). The biplot of the PCA from SST predictors (Figure S3), showed that PC2 separated populations based on the maximum SST, with CLA having a lower maximum SST compared to other sites. Additionally, for six out of the eight SNPs, the frequency of alternate alleles averaged 0.78 in the northern populations with a maximum SST of 16.4°C (CLA_1 and CLA_2). However, this average frequency dropped to 0.03 in ROS (max. SST of 19.3°C) and was below 0.01 in HLG and QUI, where the max. SST was above 21°C (Figure 4). This indicates a strong association between latitudinal change in the maximum SST and allele frequencies. The SNP ID ‘71212:99:-‘ exhibited a more gradual trend, with the alternate allele maintained at intermediate frequencies in southern populations (0.29 at ROS; 0.12 at QUI). In contrast, two outlier SNPs (‘86197:151:+’and ‘232251:239:+’, see Figure 4) showed an opposite trend, where the alternative allele reached high frequency at QUI but was absent in other populations.

**Table 5.** Summary of GLM models for eight outlier SNPs, where the model including PCs of SNP variation and SST-based PCs (ENV2 for the PC2) provided a better fit to the distribution of genotypes compared to the null model.

| Outlier ID | Model | Predictor | AIC | Pseudo $R^2$ | LL ratio test |
| --- | --- | --- | --- | --- | --- |
| 144563:157:- | NULL | SNP ~ PC1+PC4 | 40.66 | 0.608 | <0.001 |
| 144563:157:- | ENV | SNP ~ PC4+ENV2 | 39.306 | 0.623 |  |
| 174791:33:+ | NULL | SNP ~ PC1+PC5 | 66.74 | 0.607 | 0.0027 |
| 174791:33:+ | ENV | SNP ~ PC3+PC4+PC5+ENV2 | 58.919 | 0.684 |  |
| 181069:233:+ | NULL | SNP ~ PC1+PC3 | 84.622 | 0.559 | <0.001 |
| 181069:233:+ | ENV | SNP ~ PC2+PC4+ENV2 | 74.502 | 0.627 |  |
| 232251:239:+ | NULL | SNP ~ PC1+PC2 | 81.927 | 0.419 | 0.0015 |
| 232251:239:+ | ENV | SNP ~ PC1+PC2+ENV2 | 73.877 | 0.496 |  |
| 253698:139:- | NULL | SNP ~ PC1+PC5 | 64.723 | 0.69 | <0.001 |
| 253698:139:- | ENV | SNP ~ PC1+PC4+PC5+ENV2 | 50.795 | 0.784 |  |
| 35776:116:+ | NULL | SNP ~ PC1+PC3+PC4 | 60.8 | 0.715 | <0.001 |
| 35776:116:+ | ENV | SNP ~ PC1+PC3+ENV2 | 56.772 | 0.737 |  |
| 71212:99:- | NULL | SNP ~ PC1+PC3+PC4+PC5 | 68.509 | 0.719 | <0.001 |
| 71212:99:- | ENV | SNP ~ PC4+PC5+ENV2 | 50.852 | 0.794 |  |
| 86197:151:+ | NULL | SNP ~ PC1+PC2 | 56.849 | 0.73 | <0.001 |
| 86197:151:+ | ENV | SNP ~ PC2+PC3+PC5+ENV2 | 39.61 | 0.843 |  |

**Figure 4.**
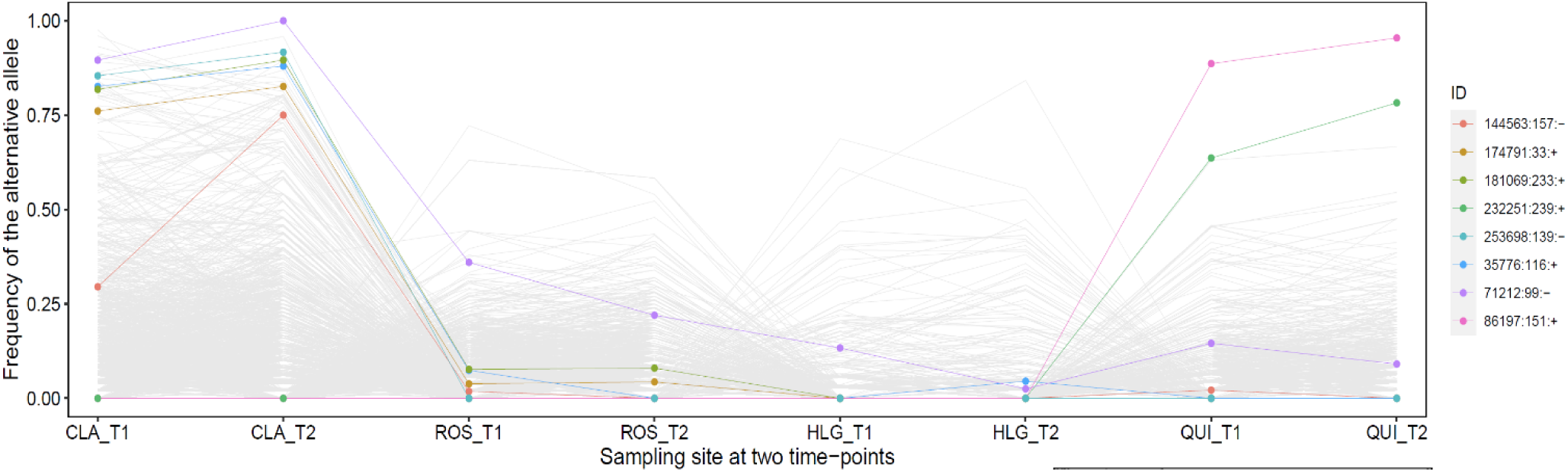
Minor allele frequencies of SNPs across space for both time points (T1 or T2) are displayed. The sampling sites are sorted in descending order based on the maximum SST. Outlier SNPs, which were identified through spatial differentiation and had genotypes that were best fitted by the ENV2 model incorporating the PC2 of the SST predictors, are color-coded according to the legend. Putatively neutral SNPs and other outliers not explained by the maximum SST are represented in grey.

### 3.7 Gene ontology

Functional analyses using the OmicsBox 2.2.4/Blast2GO pipeline were restricted to 53 loci that had SNPs detected as outliers in both space (16 SNPs) and time (38 SNPs) by at least two methods. Out of the 53 loci, nine (16.58%) showed significant hits in the BLASTX search against the UniProt database. The top hits were found to be related to brown algae of the Ectocarpus genus, with seven top hits having an e-value equal to or lower than 1.52E-4 (Table S3). Among these sequences, four sequences had SNPs detected by outlier tests across space while the remaining five sequences were detected over time. Gene Ontology functional annotation of these contigs resulted in hits for eight out of the nine significant BLASTX matches. The annotated sequences were functionally categorized into various categories, including metabolic processes, cellular processes, biological regulation, response to stimuli, and others. (Table S3).

## 4. DISCUSSION

Our study has yielded valuable insights into the contemporary evolution of populations of the kelp *Laminaria digitata* by examining spatial and temporal genetic variation. While initial genome scans conducted at a single time point indicated the potential involvement of temperature in the populations’ adaptive response, temporal genomics has provided valuable insight for deciphering the effects of ongoing selection relative to genetic drift in allele frequency changes. Below we outline two contrasting evolutionary scenarios and discuss their implication for the long-term survival of the species.

### 4.1 Signals of adaptive response at the southern margin

This study supports the theoretical expectation whereby populations at the southern margin are genetically impoverished relative to core populations due to smaller effective population size and/or geographical isolation (Blows & Hoffmann, 1993; Johannesson & André, 2006; Eckert et al., 2008; Pilczynska et al., 2019). The observed reduction in the levels of genetic diversity at the southern-most limit (QUI, Quiberon) compared to a central population (ROS, Roscoff) is consistent with previous studies using microsatellite markers (Oppliger et al., 2014; Robuchon et al., 2014; Liesner et al., 2020).

However, despite the moderately effective population size in Quiberon relative to Roscoff, there was no evidence of reduced adaptive potential in Quiberon. This is supported by the detection of, (i) candidate loci with high spatial differentiation potentially associated with maximum sea surface temperature (SST), and (ii) candidate loci with higher temporal differentiation than expected under genetic drift alone, which were reported in equivalent proportion between Roscoff and Quiberon. In addition, some of these candidate loci have important metabolic and cellular functions that could be involved in adaptive responses. While our findings support the hypothesis that SST is driving the adaptive response of *L. digitata*, consistent with previous heat stress experiments performed on the same populations (Liesner et al., 2020; Schimpf et al., 2022) and genome scans of other kelp species (Guzinski et al., 2020; Mao et al., 2020; Vranken et al., 2021), it is important to note that confounding factors may also be at play. This is particularly relevant in the present study because of the phylogeographical history of the species along the North Eastern coast of the Atlantic which indicated that *L. digitata* is composed of two different clusters (Neiva et al. 2020). The southern Europe cluster extends from southern Brittany, along the Channel and North Sea to its northern limit located in Helgoland (HLG), while the northern Europe cluster, spread from Ireland to Iceland and north Norway regions (Leisner et al., 2020; Neiva et al., 2020). Two different refugia likely contributed independently to the post-glacial colonization of the Brittany/North Sea and northern Europe regions, with one located along the Armorican coastlines and the other in Northern UK (Neiva et al., 2020). The clear genetic discontinuity suggests that genetic structuring across the older Brittany region dates back to the back to Last Glacial Maximum. This evolutionary pattern has been found in many different seaweeds of the North Eastern Atlantic, as reviewed by Neiva et al. (2016), indicating that genetic structure in seaweeds, characterized by low dispersal distances, can arise from historical processes and be maintained at relatively small spatial scales. Due to this evolutionary pattern, distinguishing the effects of selection and demography from population genetic differentiation detected through genome scans (Narum & Hess, 2011; Bierne et al., 2013; Bank et al., 2014) can be challenging. However, temporal genomics, which involves studying changes in genetic variation over time, and the ability to perform outlier tests to detect unexpected patterns of allele frequency changes, provide additional support for ongoing selection at the species’ warm edge. Although temporal resolution may not be sufficient to fully test whether climate warming is the primary driver of temporal differentiation, the detection of temporal outliers in the southernmost population highlights its uniqueness and ongoing adaptive response to rapid environmental changes.

### 4.2 When genetic drift overwhelms local selection

The study of the population from the island of Helgoland in the North Sea (HLG) gave contrasting results compared to Quiberon. We had expected to find similar signals of adaptive differentiation between Quiberon and Helgoland given that both populations experience high and comparable summer SST (Derrien-Courtel et al., 2013) and exhibit slightly higher thermal tolerance than populations from the cool margins (Liesner et al., 2020). However, the population from Helgoland has the lowest levels of genome-wide genetic diversity among those analysed here, and this diversity tended to decrease over time. In addition, candidate loci with elevated temporal differentiation were rare in Helgoland, even absent after considering those detected by at least two methods of genome scans. Together, these findings support that genetic drift relative to selection is an important component of the contemporary evolution of this population. This is in line with earlier reports, indicating low levels of genetic diversity at microsatellites in Helgoland (Liesner et al., 2020; Fouqueau, 2021) and signals of local adaptation to temperature that became less clear when comparisons have been made with gametophytes from Helgoland rather than Quiberon (Schimpf et al., 2022). Several factors linked to the geographical isolation of the population and frequent high summer temperatures that both reduced or even inhibited the reproduction of individuals and cause severe bottlenecks (Bartsch et al., 2013) are likely to be involved in its genetic impoverishment. The prevalence of genetic drift in this last population is concordant with the considerable biomass loss of *L. digitata* that was reported between 1970 and 2005 by Pehlke and Bartsch (2008). Interestingly, genetic erosion in Helgoland seems to be not specific to *L. digitata*, as the kelp *Saccharina latissima* exhibited here its lowest level of genetic diversity as well (at both microsatellites and SNPs, Guzinski et al., 2020). The eco-evolutionary dynamics of kelp populations from Helgoland may be used more broadly for a better understanding of the adaptive responses of small and fragmented populations experiencing high rates of genetic drift (e.g., Arizmendi-Mejía et al., 2015; Crisci et al., 2017).

### 4.3 Technical considerations

As the present study was based on the genome sequencing of samples that had been stored for several years before DNA extraction and library preparation, the subsequent decreases in both the number of reads and SNPs among the oldest relative to the most recent samples were not surprising. This decrease is mainly attributable to a decline in DNA quality, which may introduce various genotyping errors (Dehasque et al., 2020). However, the strict quality control measures that we implemented, along with the high genotype correspondence between replicate samples, allowed us to select a subset of loci with high accuracy. This highlights the benefit of replicate samples in inferring evolutionary processes from genomic data sets (Reynes et al., 2021), especially those based on multiple time points (Therkildsen et al., 2013a, 2013b). Moreover, if DNA quality has indeed contributed to temporal genetic changes, we expected that most of the candidate loci would be detected at Helgoland, where the oldest samples of the study come from, rather than in populations from Roscoff and Quiberon.

### 4.4 Considering other factors than selection for temporal genetic changes

As with kelp species, *L. digitata* is characterized by haplo-diplontic life history, with macroscopic diploid sporophytes alternating with microscopic haploid gametophytes. By investigating genetic variation in the macroscopic stage, our study captures half of the biphasic life cycle of the species. Although the importance of gametophytes in the contemporary evolution of the species in response to climate change is poorly understood (see for review Veenhof et al., 2022), their potential to persist as a bank of microscopic forms (e.g., Edwards, 2000; Bartsch et al., 2013) when temperatures exceed the upper thermal thresholds of sporophytes could mitigate the negative effects of genetic drift. However, our results emphasized that despite the full replenishment of the Helgoland population from its microscopic forms (Bartsch et al., 2013), genomic variation was low and decreasing over time. In addition to the unknown gametophyte’s role in shaping evolutionary processes, we discuss below the robustness of our results concerning the assumptions we made to estimate effective population size and simulate genetic change over time from the sporophyte stage. Firstly, we assumed a two-generation sampling, whereas the observation of sporophytes with shortened life spans on the warm edges relative to cool margins, although not published for this species, has been reported for the sister species *L. hyperborea* (Bartsch et al., 2008). Consequently, the impact of genetic drift may have been previously overestimated in populations inhabiting warmer climates, resulting in a lower upper limit of neutral expectation derived from simulations of temporal genetic change. While reducing the detection threshold would increase the proportion of identified candidate loci, it would not affect those exhibiting high temporal differentiation, as there was a large discontinuity in the distribution of F_ST_ values between the latter and putatively neutral loci. Moreover, candite loci displaying elevated temporal differentiation were detected with high consistency across two different scenarios of effective population size, highlighting the persistence of the signal across various simulation parameters. Secondly, we assumed no gene flow in temporal methods for estimating effective population size (Waples, 1989; Wang, 2001; Hui & Burt, 2015) and detecting outlier loci while gene flow should be carefully evaluated in patterns of temporal genetic change (e.g., Therkildsen et al., 2013a, 2013b). However, due to the notably limited levels of gene flow and population connectivity observed in kelps (Fouqueau et al., 2023), it appears that gene flow is not a significant factor in explaining substantial allelic changes over a few generations. This was supported by the absence of significant genetic changes over time when considering the full set of SNPs and the lack of detectable first-generation migrants, which would have been readily identified through admixture proportions and PCA revealing strong spatial genetic structuring. The rarity of outlier loci relative to neutral ones strengthens the hypothesis that they undergo evolutionary processes distinct from the background genomic variation. Hence, ongoing selection was considered the most plausible explanation for the identification of candidate loci exhibiting higher temporal differentiation than expected under genetic drift alone. Further studies should now clarify how strong local selection acting on both the gametophyte and sporophyte stages can lead to pronounced temporal F_ST_ within a few generations.

## Acknowledgements

This work was funded by the EU project MARFOR Biodiversa/004/2015. The authors thank the ABiMS platform of the Roscoff biological station (http://abims.sb-roscoff.fr) for providing the HPC resources that contributed to the search results reported in this document.

## Supplementary section

### Supplementary material S1

This supplementary section aims to evaluate the impact of choosing low or high Ne scenarios in the simulation framework. We performed simulations of SNP frequencies 5000 times, using the same parameters described in the “Outlier detection across time” section of the main manuscript but considering the lowest Ne estimates obtained from temporal methods (see Table 3). We expected that decreasing Ne would strengthen genetic drift, leading to increased levels of temporal neutral genetic differentiation and a higher detection threshold for outlier SNPs. However, we found no significant differences in mean F_ST_ values when comparing simulations based on the low and high Ne scenarios. The mean F_ST_ values ranged from 0.002 to 0.003 for the high and low Ne scenarios, respectively. Additionally, the detection of outlier SNPs was minimally influenced by this parameter. Person’s correlations between SNP p-values in the low and high Ne scenarios were high, with correlations of r = 0.962 at HLG, r = 0.997 at ROS and r = 0.986 at QUI. This indicates that the SNPs detected as outliers were almost identical across the two different Ne scenarios. Interestingly, reducing Ne at ROS from 43 063 to 247 prevented the detection of two outliers that were reported from the high Ne scenario. It is noteworthy that major differences were expected at ROS, considering the reduction in Ne from thousands to hundreds of individuals, while the range was narrower at HLG and QUI (around two times). This suggests that changes in allele frequencies under genetic drift over two generations may be buffered at ROS by maintaining Ne in at least two hundred individuals. However, subsequent analyses indicated that reducing Ne had a greater impact on simulations and genetic changes when genetic drift was simulated over a thousand generations, rather than two (data not shown).

**Figure S1.**
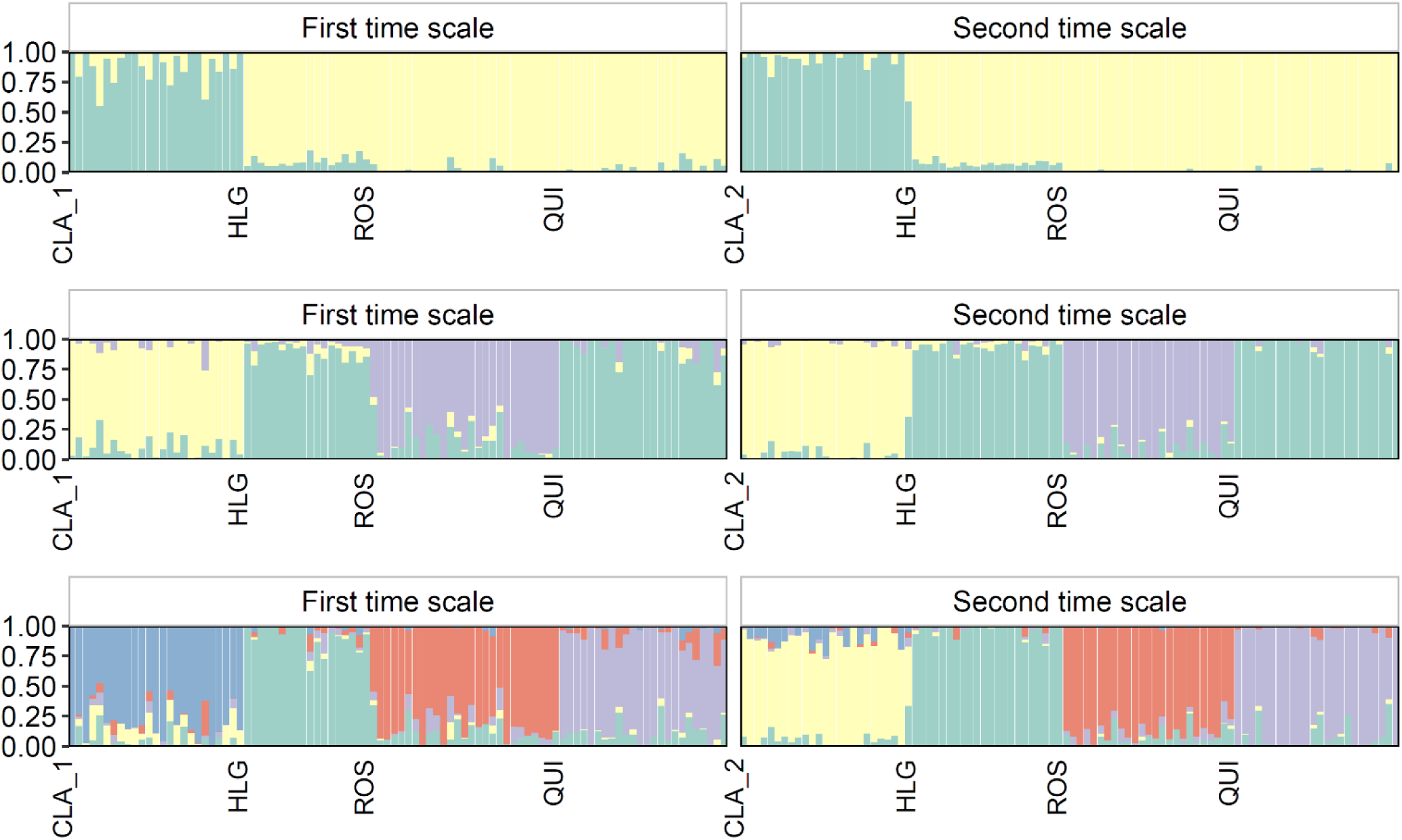
SNMF admixture plots for different values of K, including K = 2, K = 3, and the optimal number of clusters K = 5. Each bar in the plots represents one sampled individual, and the colors within the bar represent the membership proportions of each cluster.

**Figure S2.**
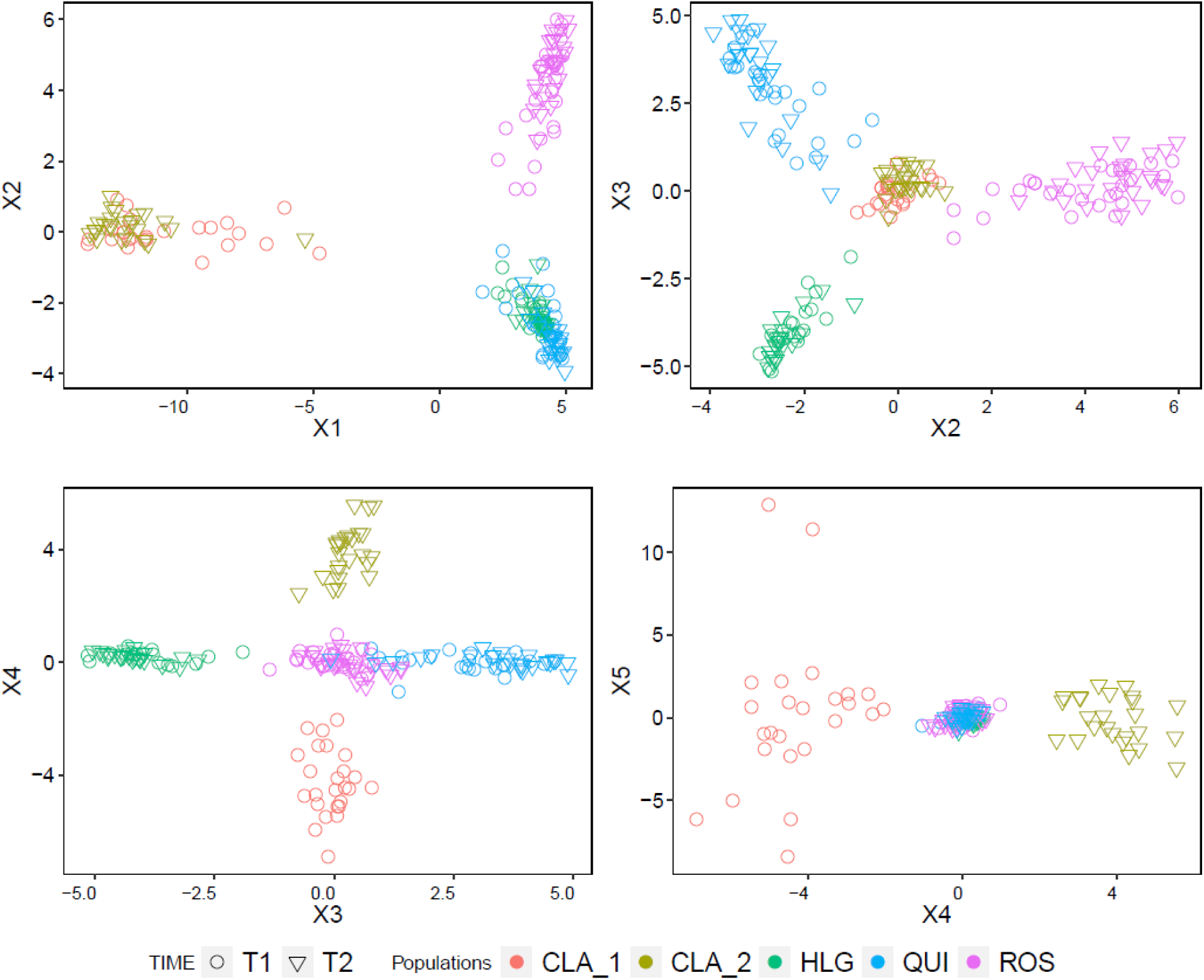
PCA biplots of SNP variation, including the five principal components: PC1 (46.5%) vs PC2 (8.3%); PC2 vs PC3 (6.1%), PC3 vs PC4 (4.5%) and PC4 vs PC5 (2.7%).

**Figure S3.**
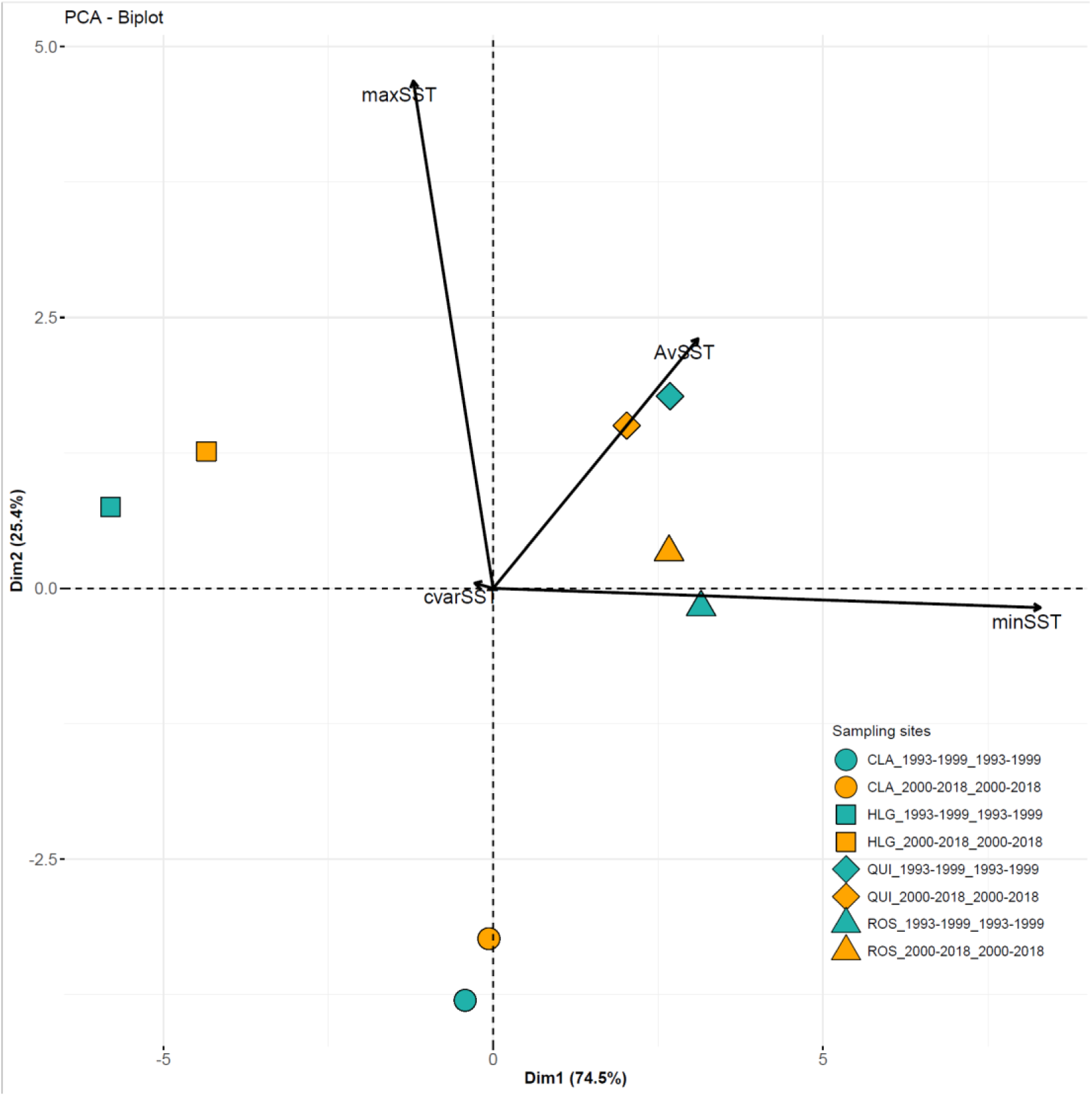
Biplot of the first two PCs derived from the PCA conducted on daily mean SST statistics. The positions of samples represent their scores on the first two PCs, indicating their similarity or dissimilarity in terms of SST variation. Both PCs were included as explanatory variables in generalized linear models (GLMs) to assess the influence of local genetic drift and temperature on the distribution of genotypes at focal outlier SNPs.

**Table S1.** Individual missingness, and mean read depth (Mean DP) were assessed at both time points. Individual missingness refers to the average proportion of missing genotypes for each individual. Mean read depth was calculated by averaging the read depth across all sites. The SNP error rate estimated using eight replicates is also indicated.

| Population | Time point | n | Ind. missingness | Mean DP | SNP error rate |
| --- | --- | --- | --- | --- | --- |
| HLG | 1st | 18 | 0.05 | 13.218 | 0.020 |
|  | 2nd | 22 | 0.02 | 11.640 | 0.012 |
| CLA | 1st | 25 | 0.04 | 12.497 | 0.036 |
|  | 2nd | 25 | 0.02 | 26.362 | 0.002 |
| ROS | 1st | 27 | 0.03 | 28.457 | 0.021 |
|  | 2nd | 25 | 0.02 | 26.177 | 0.001 |
| QUI | 1st | 24 | 0.06 | 10.570 | 0.007 |
|  | 2nd | 24 | 0.02 | 24.703 | 0 |

**Table S2.** Pairwise F_ST_ values between populations and the comparisons between time points are highlighted in bold in the lower diagonal matrix. The p-values of the genic differentiation tests are displayed in the upper diagonal matrix.

| $F_{ST}$ | HLG (T1) | HLG (T2) | CLA_1 | CLA_2 | ROS (T1) | ROS (T2) | QUI (T1) | QUI (T2) |
| --- | --- | --- | --- | --- | --- | --- | --- | --- |
| HLG (T1) |  | **NS** | < 0.001 | < 0.001 | < 0.001 | < 0.001 | < 0.001 | < 0.001 |
| HLG (T2) | **0.008** |  | < 0.001 | < 0.001 | < 0.001 | < 0.001 | < 0.001 | < 0.001 |
| CLA_1 | 0.257 | 0.285 |  | **< 0.001** | < 0.001 | < 0.001 | < 0.001 | < 0.001 |
| CLA_2 | 0.266 | 0.297 | **0.039** |  | < 0.001 | < 0.001 | < 0.001 | < 0.001 |
| ROS (T1) | 0.094 | 0.113 | 0.224 | 0.235 |  | **NS** | < 0.001 | < 0.001 |
| ROS (T2) | 0.104 | 0.125 | 0.223 | 0.237 | **0.000** |  | < 0.001 | < 0.001 |
| QUI (T1) | 0.127 | 0.150 | 0.251 | 0.262 | 0.078 | 0.083 |  | **NS** |
| QUI (T2) | 0.142 | 0.167 | 0.249 | 0.272 | 0.090 | 0.095 | **0.003** |  |

**Table S3.**
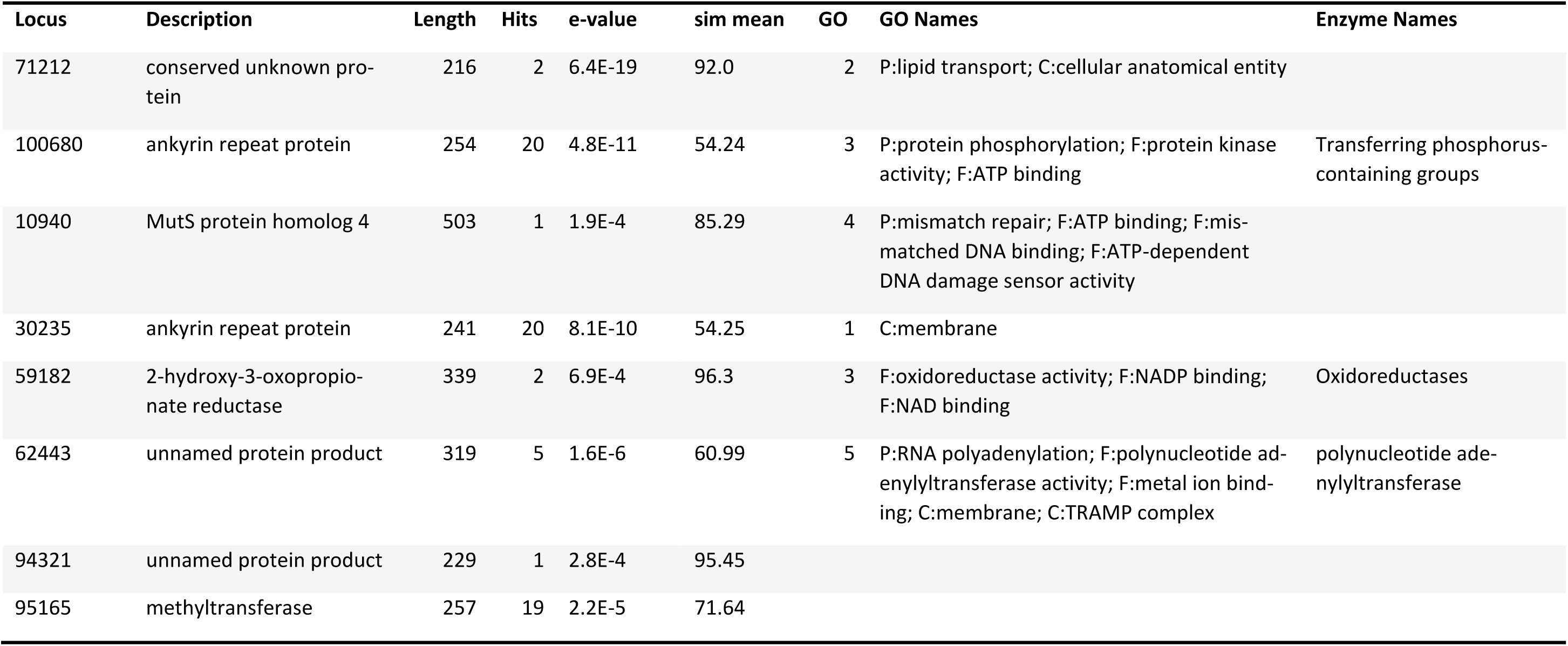
Functional categorization of eight annotated loci derived from outlier SNPs out of the 53 loci.

| Locus | Description | Length | Hits | e-value | sim mean | GO | GO Names | Enzyme Names |
| --- | --- | --- | --- | --- | --- | --- | --- | --- |
| 71212 | conserved unknown protein | 216 | 2 | 6.4E-19 | 92.0 | 2 | P:lipid transport; C:cellular anatomical entity |  |
| 100680 | ankyrin repeat protein | 254 | 20 | 4.8E-11 | 54.24 | 3 | P:protein phosphorylation; F:protein kinase activity; F:ATP binding | Transferring phosphorus-containing groups |
| 10940 | MutS protein homolog 4 | 503 | 1 | 1.9E-4 | 85.29 | 4 | P:mismatch repair; F:ATP binding; F:mismatched DNA binding; F:ATP-dependent DNA damage sensor activity |  |
| 30235 | ankyrin repeat protein | 241 | 20 | 8.1E-10 | 54.25 | 1 | C:membrane |  |
| 59182 | 2-hydroxy-3-oxopropionate reductase | 339 | 2 | 6.9E-4 | 96.3 | 3 | F:oxidoreductase activity; F:NADP binding; F:NAD binding | Oxidoreductases |
| 62443 | unnamed protein product | 319 | 5 | 1.6E-6 | 60.99 | 5 | P:RNA polyadenylation; F:polynucleotide adenylyltransferase activity; F:metal ion binding; C:membrane; C:TRAMP complex | polynucleotide adenylyltransferase |
| 94321 | unnamed protein product | 229 | 1 | 2.8E-4 | 95.45 |  |  |  |
| 95165 | methyltransferase | 257 | 19 | 2.2E-5 | 71.64 |  |  |  |

